# A Kuramoto model of self-other integration across interpersonal synchronization strategies

**DOI:** 10.1101/640383

**Authors:** O.A. Heggli, J. Cabral, I. Konvalinka, P. Vuust, M.L. Kringelbach

## Abstract

Human social behaviour is complex, and the biological and neural mechanisms underpinning it remain debated^1,2^. A particularly interesting social phenomenon is our ability and tendency to fall into synchrony with other humans^3,4^. Our ability to coordinate actions and goals relies on the ability to distinguish between and integrate self and other, which when impaired can lead to devastating consequences. Interpersonal synchronization has been a widely used framework for studying action coordination and self-other integration, showing that in simple interactions, such as joint finger tapping, complex interpersonal dynamics emerge. Here we propose a computational model of self-other integration via within- and between-person action-perception links, implemented as a simple Kuramoto model with four oscillators. The model abstracts each member of a dyad as a unit consisting of two connected oscillators, representing intrinsic processes of perception and action. By fitting this model to data from two separate experiments we show that interpersonal synchronization strategies rely on the relationship between within- and between-unit coupling. Specifically, *mutual adaptation* exhibits a higher between-unit coupling than within-unit coupling; *leading-following* requires that the follower unit has a low within-unit coupling; and *leading-leading* occurs when two units jointly exhibit a low between-unit coupling. These findings are consistent with the theory of interpersonal synchronization emerging through self-other integration mediated by processes of action-perception coupling^4^. Hence, our results show that chaotic human behaviour occurring on a millisecond scale can be modelled using coupled oscillators.

## Background

When two people perform a simple task together, such as walking together or applauding a successful performance, they tend towards synchronization^5,6^. This emergence of synchrony is also found in many other natural phenomena^3^, such as the collective flashings of fireflies^7^, or the pacemaker cells in the heart^8^. For this reason, the mathematical framework of coupled oscillators provides an approach for understanding the conditions and parameters necessary for synchrony to emerge^9–11^. However, in many cases of human interaction, synchronization is not simply emergent, but rather a goal or a prerequisite of the task. A particularly prominent example of this is rhythmic joint action, as found in musical performance. Here, multiple people coordinate their movements and adapt to each other on a millisecond basis. In this sense, human interpersonal synchronization presents as a more complex system. Experiments using joint finger tapping paradigms (illustrated in figure 1a and 1b) shows that this type of synchronization relies on different synchronization strategies, such as mutual adaptation and leading-following^4,12–14^. Common for these is that they cannot be differentiated by looking just at measures of synchronization, as different strategies may exhibit the same synchronization level. Instead, differences between synchronization strategies can be detected when a lagged cross-correlation is calculated between the resultant time-series, as illustrated in figure 1C and 1D^12^.

**Figure 1.**
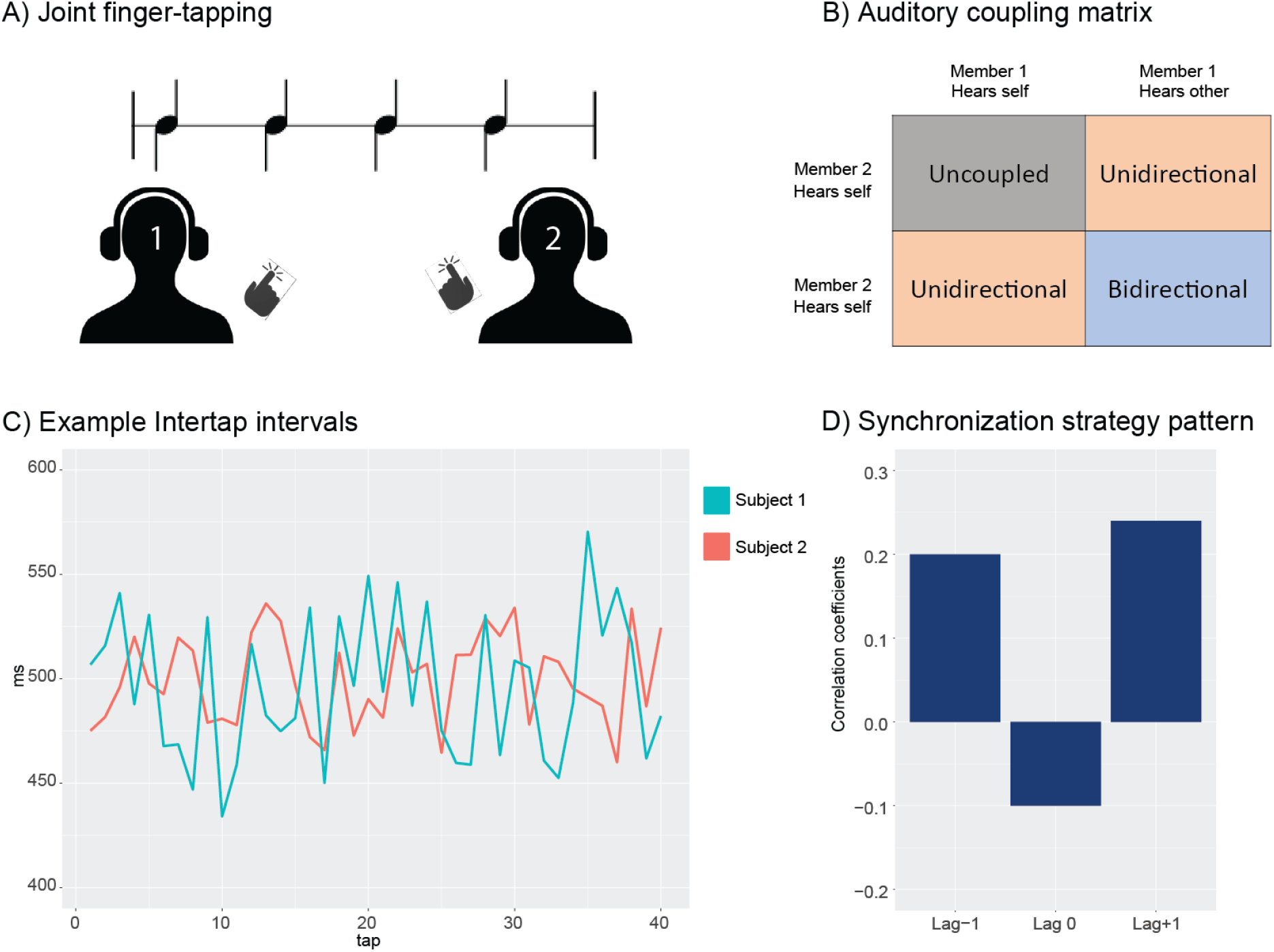
In A) a joint finger tapping paradigm is illustrated. Two persons (dyad members) tap an isochronous rhythm together. Their auditory feedback is shown in matrix form in B). C) Time series representing the intertap interval (ITI), a measure of the time between successive taps, of each dyad member. Colours indicate dyad member. When these time series are cross-correlated at lag −1, lag 0, and lag +1, a pattern such as illustrated in D) emerges. Here, the pattern would indicate a mutual adaptation synchronization strategy.

The most commonly found strategy is the one of mutual adaptation, which occurs when both members in an interacting dyad simultaneously and constantly adapt to each other on a per-action basis^4,15^. This results in positive correlations at lag −1 and lag +1, and a negative correlation at lag 0, as the members are mutually correlated with the previous tap of the other. Another well-documented synchronization strategy is the leading-following strategy, where one of the dyad members exhibits less adaptability than the other and hence becomes a leader. This shows as a positive correlation at either lag +1 or lag −1, depending on which member is leading. While both these strategies have been reported in multiple studies, recently a third strategy called leading-leading was found^5,12,13,16^. In the leading-leading strategy both members resist adaptation and rather taps along without much regard to the performance of their tapping partner. This results in a pattern with low correlation coefficients across all lags. While there have been attempts at modelling such behaviour with coupled oscillators the existing models have not captured the mechanisms underlying these distinct synchronization strategies^5,17–20^.

One explanation for the emergence of these different strategies may lie in the increased complexity compared to physical systems such as coupled pendulums. In humans, the coupling is mediated by perceptual links, such as visual or auditory information, as opposed to the physical coupling found in many other systems^4^. Unsurprisingly, if no such link is present (for instance if no perception of the other’s movement is possible) experiments show that synchronization does not occur^21^. Hence, interpersonal synchronization necessitates two separate processes, wherein one process is perceiving the stimuli to be synchronized to, and another process is in charge of producing the actions leading to synchronization. The last couple of decades of research points towards these two processes, action and perception, being intrinsically coupled in terms of processing in the human brain^22,23^. In joint finger tapping such action-perception coupling can occur when one dyad member perceives the auditory feedback from the other member as belonging to its own tapping, hence blurring the lines between self and other. It is this type of coupling that has been hypothesized to underlie the mutual adaptation synchronization strategy observed in previous studies^4^. On the other hand, if one dyad member chooses to ignore feedback from the other dyad member and instead solely monitors their own model of the task, this will force the other dyad member to take over the coordination task, and thus create a leading-following relationship. By necessity, this then requires the leading dyad member to decouple their motor actions from their auditory perception of the other.

Here we test the hypothesis that synchronization strategies emerge as a function of action-perception coupling strength. Specifically, we test if the rhythmic joint finger tapping tasks commonly used in the field of joint action can be modelled using a coupled oscillator model, and if empirically encountered synchronization strategies are systematically linked to differing coupling strengths in the model. In the first part, we examine the behaviour of a four-oscillator Kuramoto model to determine if it is able to reach and maintain synchronization within the short amount of time as is seen in joint finger tapping experiments, while at the same time producing distinct synchronization strategies. In the second part we use empirical data to validate the model, and determine how coupling parameters are linked to specific synchronization strategies.

## Model of interacting dyads

Our model aims at representing the dynamics of interacting dyads performing joint finger tapping in a reduced form, retaining only the necessary features to capture the fundamental principles underlying the complex synchronization strategies observed in joint finger tapping. Each person is considered a unit, with two internal oscillators serving as proxies for perception and action (see Figure 2A). These two within-unit oscillators are bidirectionally linked through the internal coupling term *i* representing the intrinsic coupling found between auditory and motor processes in the brain. The two units are coupled so that the action oscillator in one unit is unidirectionally linked to the perception oscillator in the other unit, through the external coupling term e. This coupling term represents the extrinsic flow of information between the two interacting units. The model is based on the Kuramoto model of coupled oscillators, with the exception that the connection strength between each pair of oscillators *n* and *p* is defined by the coupling matrix *K_np_* (Figure 2B), as defined by the following equation:

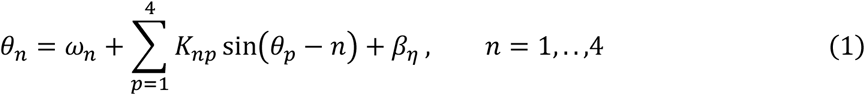

where *θ_n_* is the phase of each oscillator *n, ω_n_* is its fundamental frequency, and where *β_η_* represents added Gaussian white noise, with *η* drawn from a Gaussian distribution with standard deviation *β* (see full details in the extended Methods section).

**Figure 2.**
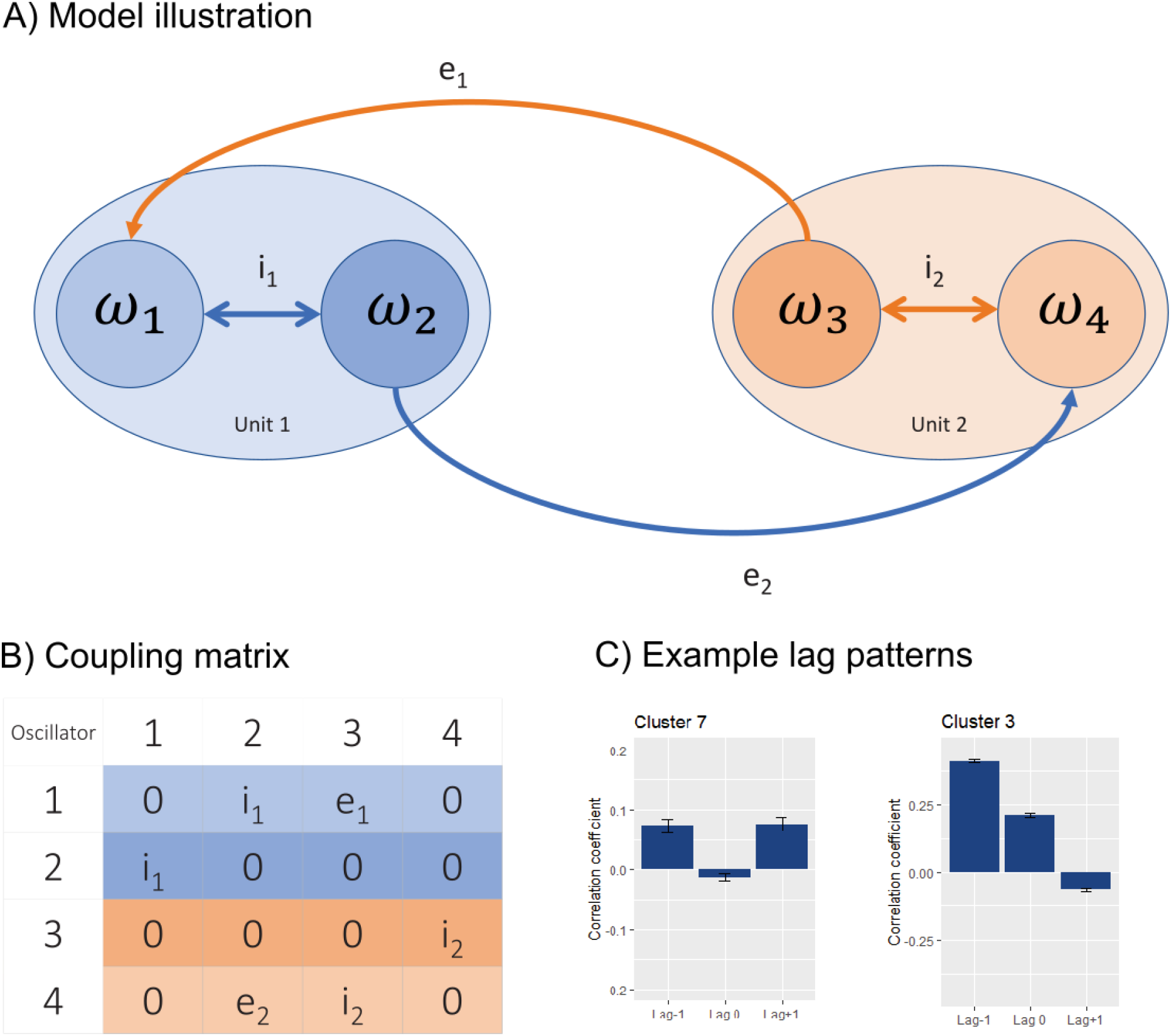
Overview of the model. In A we see the four-oscillator model, with the oscillators represented as circles within the two units. The coupling terms are showed as arrowed lines. In B the coupling matrix is shown, and two out of 13 significantly different lag patterns produce by the model is shown in C.

To examine the behaviour of the model we first determine the coupling strength producing maximum synchronization, by globally varying the four coupling terms equally (for further details see the methods section). This allowed us to determine the range of coupling strengths for which the model switches from exhibiting unsynchronized to fully synchronized behaviour. Following this, we sampled the model at a selection of coupling strength combinations, and calculated cross-correlation lag patterns of the output from each unit’s action oscillator (ω_2_ and ω_3_ in Figure 2A). These lag patterns where thereafter clustered to identify significantly different lag patterns.

### Results

Simulations showed that the model reached a maximum synchronous state at a general coupling weight of 15.5, as measured with the synchronization index^24^. This index is calculated based on the variance of relative phase between two signals, and is a unitless number ranging from 0 to 1, with 1 indicating full synchronization. Subsequently, we ran further simulations for a range of selected coupling weights (see methods) below this critical range to determine if the model was able to produce distinct synchronization strategies. We found that the model exhibited a rich and varied sample of lag patterns, with 13 of these patterns being different at α<0.001. Multiple clusters produced by the model have patterns with resemblances to a leading-following strategy (see Cluster 3 in figure 2C) and one cluster exhibits features of a mutual adaptation strategy (see Cluster 7 in figure 2C). Hence, we found that a model with four coupled oscillators and four coupling terms are able to produce an array of differing synchronization strategies.

To verify if this behaviour was dependent on each unit having two oscillators, we tested an even more reduced model containing only one oscillator per unit. This reduced model failed at producing lag patterns consistent with empirical data, by showing a large positive lag 0 component at all coupling strength combinations (see methods).

## Model validation on empirical data

To validate our model, we tested it on empirical data from two separate joint finger tapping studies. Dataset 1 was acquired from a 2018 study by Heggli et. al^16^, and dataset 2 was acquired from a 2010 study by Konvalinka et. al^12^. Both datasets were collected in compliance with local and national research ethics standards.

In dataset 1, musicians were paired and asked to tap one of two rhythms together while bidirectionally coupled. Both rhythms had an ITI of 500 ms, corresponding to a beat-per-minute (bpm) of 120. A cluster-analysis of the cross-correlated lags identified three subgroups of participants. One subgroup used the leading-leading strategy, and the two remaining subgroups exhibited patterns of mutual adaptation at two different pattern strengths.

In dataset 2, pairs of non-musicians tapped together in differing auditory coupling conditions: 1) uncoupled, with no auditory feedback from the other dyad member, 2) unidirectional coupling so that dyad member 1 hears their own tapping sound and member 2 hears member 1’s tapping, and mirrored so that member 2 hears its own tapping and member 1 hears member 2’s tapping, and 3) bidirectional coupling wherein dyad member 1 only hears the taps of dyad member 2 and vice versa. In addition, the tapping was performed at different tempi (96 bpm, 120 bpm, and 150 bpm). Here, a leading-following was found in the unidirectional condition, and mutual adaptation in the bidirectional condition. For the purposes of model validation, we chose only the 120-bpm tempo from this dataset, and only the conditions wherein the dyad members interacted (unidirectional coupling, and bidirectional coupling). Together, these two datasets contain three distinct synchronization strategies, in six independent groups.

We performed a two-step consecutive parameter search to determine which coupling weights best fit the data. For dataset 1 we used the three subgroups found in the empirical data, and for dataset 2 we used data from three different coupling conditions (bidirectional coupling, unidirectional coupling 1 with dyad member 1 being the leader, and unidirectional coupling 2 with dyad member 2 being the leader). We optimized our search on the numerical distance between the averaged cross-correlated lags in the empirical data and the data produced by our model. Once the best fit was found, we calculated the mean Bhattacharyya coefficient between the empirical and simulated data to serve as a measure of goodness of fit^25^.

### Results

Our parameter search provided a good fit for all subgroups found in the empirical data (see Figure 3). We found that the three empirically encountered synchronization strategies, as reproduced by our model, are characterized by the weighting of between- and within-unit couplings. Mutual adaptation, the most common synchronization strategy, relies on a high between-unit coupling strength and a correspondingly low within-unit coupling strength. Leading-following, in our case the unidirectionally forced type of leading-following, relies on the leader unit having a balanced coupling strength on both the within- and between-unit coupling term. However, the follower unit exhibits a much stronger between-unit coupling strength than its within-unit coupling. The remaining strategy, leading-leading, presents as both units having a strong within-unit coupling strength, and a low between-unit coupling strength. Overall, the mean Bhattacharyya coefficient were over 0.8 in all cases except for one instance of the mutual adaptation strategy found in in dataset 2 (see table 1).

**Figure 3.**
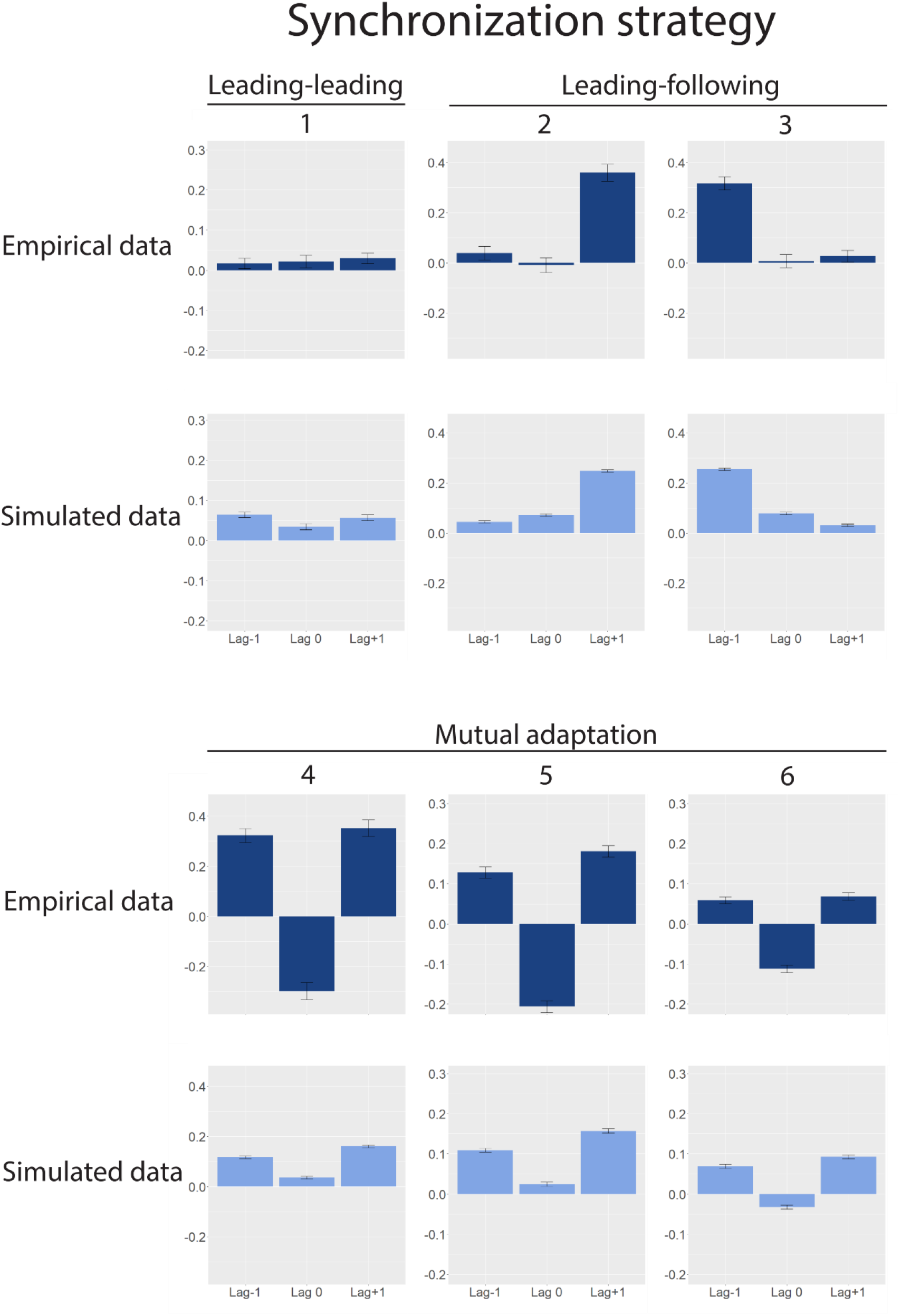
Overview of the main results. In the first row, synchronization patterns from the empirical data are shown. Leading-leading (1) corresponds to a subgroup from dataset 1. Leading-following (2) and (3) are from the two unidirectional conditions in dataset 2. From the mutual adaptation group (4) is the bidirectional condition from dataset 2, whereas (5) and 6) are the two remaining subgroups from dataset 1. The dark blue lag patterns show the empirical data. The light blue lag patterns show the synchronization strategy patterns produced by the model at the given coupling weights listed in table 1. The patterns are plotted as the mean value, with error bars indicating the standard error of the mean. Note that the y-axis is not identical in all the plots, as the strength of the pattern varies in the empirical data.

**Table 1.** Overview of the best coupling weights found for each group. The numbering of the groups corresponds to the labelling used in Figure 3. The Bhattacharyya coefficient listed here is the mean coefficient between the three lags. For details, see methods section.

| Overview of best coupling weights |  | Unit 1 |  | Unit 2 |  |
| --- | --- | --- | --- | --- | --- |
| *Coupling term* | i<sub>1</sub> | e<sub>1</sub> | i<sub>2</sub> | e<sub>2</sub> | Bhattacharyya coefficient<br>(mean) |
| 1. Dataset 1 - *Leading-leading* | 6.5 | 1.5 | 7.8 | 1.3 | .97 |
| 2. Dataset 2 - *Leading-following (1)* | 1.7 | 5.5 | 4.1 | 5.5 | .82 |
| 3. Dataset 2 - *Leading-following (2)* | 4.1 | 5.7 | 1.7 | 4.5 | .81 |
| 4. Dataset 2 - *Mutual adaptation (1)* | 2.5 | 6.3 | 2.3 | 5.1 | .77 |
| 5. Dataset 1 - *Mutual adaptation (2)* | 2.5 | 4 | 2.3 | 8 | .93 |
| 6. Dataset 1 - *Mutual adaptation (3)* | 1 | 5 | 1.3 | 3.3 | .98 |

## Discussion

Our results show that complex human behaviour can be described by a reduced model consisting of four oscillators and four coupling terms. While there have been multiple previous attempts at modelling interpersonal synchronization using either an information-processing or a dynamical systems approach, our work is the first to reproduce all three empirically-observed synchronization strategies^5,17–20^. We find that these strategies rely on the balance of within- and between-unit coupling strengths in our model, and are placed at different points in the parameter space of the model. Mutual adaptation is found when the interacting units symmetrically downregulate their within-unit coupling strength with a corresponding increase in the between-unit coupling strength. Leading-leading is found on the opposite side of this symmetric axis, with both units exhibiting a higher within-unit coupling strength than the between-unit coupling strength. We found that the leading-following synchronization strategy requires two asymmetric units, with the follower unit having a strong between-unit coupling and leading unit a balanced between- and within-unit coupling. If we consider the unit’s oscillators to represent processes of auditory perception and motor action our findings are consistent with theories positing that synchronization strategies emerge as functions of action-perception coupling^4^.

Given these results, mutual adaptation in bidirectionally coupled joint finger tapping can be seen as a form of self-other merging, whereby two interacting people collectively attribute the auditory feedback stemming from their tapping partner as intrinsically linked to their own tapping actions. This results in an interaction wherein both dyads continuously and reciprocally adapt their tapping to each other on a tap-to-tap basis. In this sense, the dyad members can be considered to actively and collectively work towards minimizing the difference between their action and the related auditory feedback, resulting in a strong interpersonal action-perception coupling. This leads to the dynamics of the oscillators being predominantly governed by the information flow between the units, as expected in mutual adaptation. In the leading-leading strategy, this relationship is reversed.

The most likely behavioural explanation of leading-leading is that the dyad members both decouple the self-other loop, and instead focus on their own representation of the task^16^. Accordingly, our best fit for this strategy shows that both units exhibit a strong within-unit coupling, and a weak between-unit coupling. Hence, information flowing between the two units does not, to a noticeable degree, impact the behaviour of the individual unit. A key factor in understanding the emergence of this synchronization strategy is that in the behavioural experiment is that the two participants are in fact strongly coupled to the same external metronome at the start of the task. In our model, this is reflected in the starting frequency of the oscillators, and given a strong enough within-unit coupling this frequency is preserved to the point where synchronization occur without the need for a strong between-unit coupling to modulate any deviance in the starting frequency. Hence, given two participants with sufficient skill it is possible to exhibit synchronization, predominantly as an artefact of their beat-keeping skills. However, the small, but non-zero, between-unit coupling may then function as an error-detection threshold, such that if one participant should strongly deviate the other may still choose to follow.

For the leading-following strategy wherein the leader hears themselves and the follower only hears the leader, the model converges on the leading unit having a balanced weight on its within- and between-unit coupling. The follower unit exhibits a stronger self-other (between-unit) coupling than its within-unit coupling. To achieve synchrony, the follower needs to consistently monitor the auditory feedback coming from the leader, while the leader is decoupled from the follower. It is interesting that the model here converges to a balanced within- and between-unit coupling in the leading unit. One likely explanation for this comes from the characteristics of the participants in dataset 2, which were all non-musicians. This is evident in the increased noisiness of their tapping, and we reflected this in the simulations by having an increased noise level as compared to the simulations for dataset 1 (see methods). Having a balanced within-unit coupling may then act as a sort of self-correcting behaviour, ensuring that the contribution from noise in the individual oscillator is kept from increasing to heavily. It should also be noted that the leading-following behaviour seen here is experimentally forced due to the unidirectional auditory coupling between the participants. Previous research has shown that the leading-following strategy can also occur in cases of bidirectional auditory coupling, and that leaders can be distinguished from followers by increased frontal alpha suppression measured with EEG^14^. This finding has been interpreted to indicate an increase in cognitive load for leaders in bidirectionally coupled leading-following, due to the need for separating the auditory information from the followers from their own tapping actions. In this case, it may be that unidirectional leading-following differs from bidirectional leading-following, and it remains a possibility that our model would choose differing coupling weights for this strategy dependent on the participant’s auditory feedback.

The weakest point in our model appears to be the lag 0 component in the mutual adaptation strategy, as is evident from the low measure of fit found in the bidirectional condition in dataset 2. A likely explanation for this is that our model is continuously coupled, and is therefore able to adjust also when there would be no information present in a real-world setting, such as between taps. A solution for this would be to couple the two units intermittently, so that the coupling only exists when one unit produces an output. In addition, due to limitations in the perceptual threshold, we would suggest including a filtering method wherein such information would only be passed between the units if it exceeds a pre-defined tolerance region, akin to mechanisms of predictive coding^26^. For future work we would therefore suggest that incorporating time varying coupling weights could prove beneficial towards modelling human behaviour with coupled oscillators.

As we have discussed above, all three synchronization strategies can be interpreted as emerging from different combinations and strengths of within- and between-person action-perception links. A likely explanation for how and why such action-perception links emerge can be found in mechanisms of self-other integration. There is ample evidence that the human brain processes perceived and performed actions using overlapping networks (for a review see Keysers and Gazzola 2009)^27^. For instance, observing an action can produce activity in motor areas of the brain, and observing someone else being touched can lead to activity in somatosensory regions in the brain^28^. Hence, there needs to be a mechanism that distinguishes between actions related to the self, and to others, commonly referred to as self-other representation. This refers to the process of categorizing whether a percept belongs to self- or other-produced actions. One way of considering action-perception coupling would then be that it occurs as the result of minimizing the distance between self- and other-representations. In other words, in tapping tasks such as those used in this work, action-perception coupling may stem from participants categorizing the auditory feedback they hear as related to their own tapping actions, instead of belong to their tapping partner. This view of action-perception coupling finds support in the brain’s tendency towards minimizing computing costs, as formalized by Friston’s work on the free energy principle^4,29–31^. Here, the brain is considered to constantly strive for energy optimized representations of its environment. In joint finger tapping, minimizing the difference in self-other representation decreases the need for maintaining a cognitive model of the tapping partner’s behaviour. However, while this would mean that mutual adaptation is a strong attractor state, it is not the only stable synchronization strategy in interpersonal synchronization as both leading-following and leading-leading have been shown to emerge in cases of bidirectional coupling.

As with any computational model of a complex real-world process, our model can only approximate the processes involved. When considering the neural underpinnings of interpersonal synchronization, our model does not make any strict assumption. Rather, the structure of the model may be interpreted to represent action-perception coupling, as part of the more complex processes of self-other representation. Likely, these processes are all involved in interpersonal synchronization and in the selection of synchronization strategies. As rhythmic interpersonal synchronization requires auditory perception, and motor action, we expect brain regions and networks linked to such to be involved. It is also likely that regions and networks linked to social cognition is involved. For instance, our data includes a case of bidirectional coupling resulting in two distinctly different synchronization strategies, mutual adaptation and leading-leading. Previous research has also shown the existence of leading-leading in cases of bidirectional auditory coupling^14^. Hence, there needs to be a neural structure or network involved in the selection of synchronization strategy, that is independent of auditory coupling. A likely candidate here is the temporoparietal junction (TPJ). This region, located where the temporal and parietal lobes meet, has been shown to act as a network node between the thalamus and the limbic system, as well as with sensory systems^32^. In particular, the right TPJ are involved in segregating self-produced actions from actions produced by others, as is shown in lesion studies and in studies using transcranial magnetic stimulation^33,34^. We would therefore hypothesize that the involvement of the TPJ, either as a separate region or as part of a distributed network, is a key factor in the emergence of synchronization strategies. An interpretation of our model is then that the two oscillators represents an interplay between auditory and motor regions mediated by the TPJ, as shown in figure 4. Here, the between-brain couplings rely on the auditory perception of the motor actions produced by the other. We hypothesize that synchronization strategies may be distinguished in electrophysiological recordings by activity in such a network. For instance, mutual adaptation likely requires neural synchronization between representation of self- and other, as mediated in tapping tasks by motor and auditory systems in the brain^4^. Hence, one would expect to see more coherent activity between the involved brain regions during mutual adaptation than in leading-leading or leading-following.

**Figure 4.**
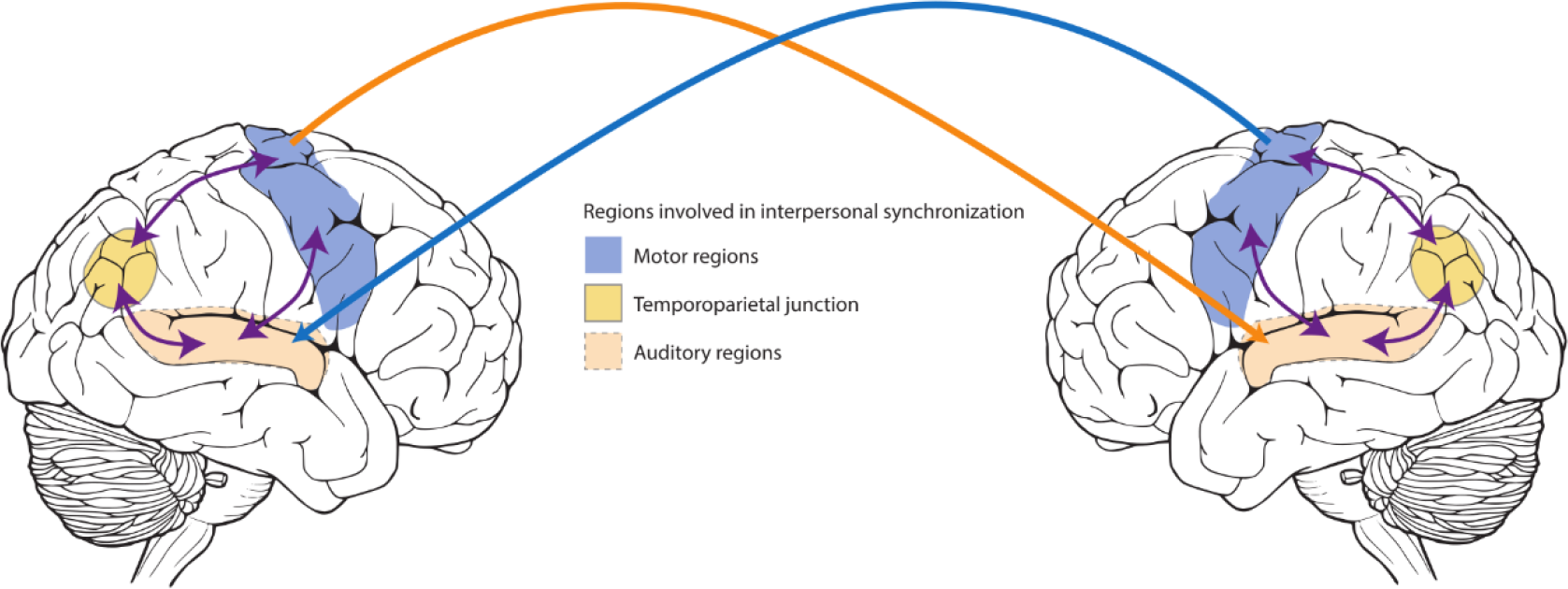
Illustration of regions involved in interpersonal synchronization. Motor regions (shown in blue) are bidirectionally linked to auditory regions (shown in light red), and with the temporoparietal junction (shown in yellow). Actions produced by one individual’s motor system is perceived in auditory regions of the other individual.

**Figure 5.**
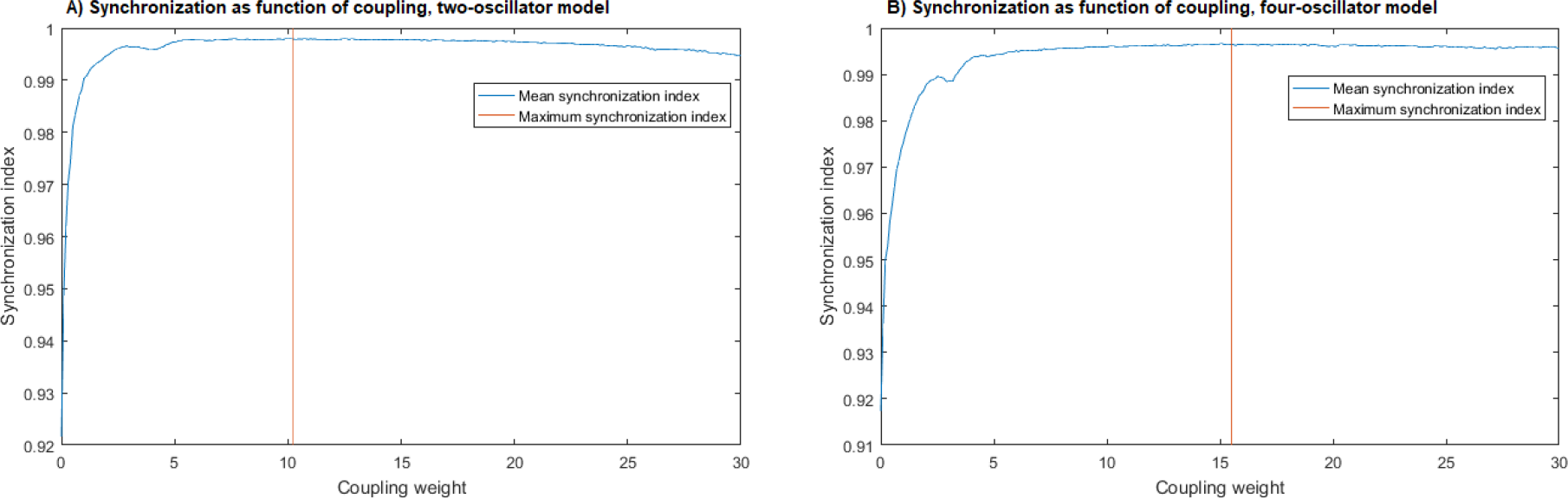
Synchronization as measured by the synchronization index as a function of coupling weights. In A we see the synchronization index of the two-oscillator model as a function of coupling weight. The vertical orange indicates the point of maximum synchronization. In B the same is shown for the four-oscillator model.

## Conclusion

In this paper we have shown how synchronization strategies found in human joint action may be successfully modelled using a reduced model consisting of four oscillators and four coupling terms. We found that synchronization strategies can be distinguished based on their within- and between-unit coupling strengths. For this particular study we interpret our model within the framework of self-other integration via within- and between-person action-perception links. However, we believe the model may be successfully applied to many other types of behaviour, such as modelling groups of people and other processes relying on perceptually mediated couplings between individuals. The model is easily scalable, although increasing the interacting units comes at computational cost. In informal tests with up to 200 interacting units we observe complex behaviours such as short-lived stable states of both in- and anti-phase synchronization. Hence, the model may be a promising tool for exploring network topologies in multi-person interactions, such as for instance in symphony orchestras. We furthermore present a likely neural interpretation of the model, where we suggest that interpersonal synchronization strategies may be represented as coherence in a network between auditory and motor regions, and the temporoparietal junction. Our model and its behaviour suggest that complex human behaviour may be the result of simple interacting components, and that coupled oscillators are able to capture these dynamics.

## Author contributions

OAH: Conceptualization, Software, Data curation, Methodology, Analysis, Validation, Visualization, Writing - original draft, Writing – review & editing.

JC: Conceptualization, Software, Writing – review & editing.

IK: Data curation, Analysis, Validation, Writing – review & editing.

PV: Funding acquisition, Analysis, Writing – review & editing.

MLK: Conceptualization, Methodology, Software, Visualization, Writing – review & editing.

## Data Availability Statement

The model implementation used in this manuscript is under active development and available on https://github.com/OleAd/FourOscModel. The behavioural data in dataset 1 is available upon request from Ole Adrian Heggli. The behavioural data in dataset 2 is available upon request from Ivana Konvalinka.

## Acknowledgments

OAH, PV and MLK is supported as part of the Center for Music in the Brain, Danish National Research Foundation (DNRF117).

MLK is supported by the ERC Consolidator Grant: CAREGIVING (615539).

JC is supported under the project NORTE-01-0145-FEDER-000023 from the Northern Portugal Regional Operational Program (NORTE2020).

IK is supported by The VILLUM Experiment grant, “Enhancing social interaction through real-time two-brain imaging”.

## Methods

### Implementation of the models

We implemented the models in MATLAB R2016b^35^, using a script for performing numerical integration of the Kuramoto model of coupled oscillators^36,37^. The oscillators are described in Equation 2 and are based on the Kuramoto model, with the exception that the between-oscillator couplings are defined by the matrix *K_ij_*. For the two-oscillator model we used two coupling terms as shown in Equation 3. For the four-oscillator model we used four coupling terms in six couplings as shown in Equation 4. Both models have a gaussian noise component *β_η_* with *η* drawn from a Gaussian distribution with standard deviation *β.* This was included to account for natural variability and noisiness observed in empirical data. For study 1 and dataset 1 in study 2 the standard deviation was set to be equivalent to 20 ms, corresponding to the interquartile range (IQR) of the inter-tap intervals (ITIs) observed in the empirical data used for study 2^16^. For dataset 2 in study 2, the standard deviation of the noise was set to an equivalent to 34.5 ms, corresponding to the IQR of the ITIs in dataset 2. In all simulations the oscillators intrinsic frequency *ω* was set to 2 Hz, with a standard deviation of 0.2 Hz, to account for natural variations in the frequency locking ability of the participants. The oscillators were initiated at random phases. For inspecting the model’s performance and behaviour, we calculated the phases of the oscillators in steps of 25ms and sampled the phase at intervals of 500 ms for computational efficiency, and linearly interpolated the zero-crossing point as a basis for creating a time-series of tap events. From this time-series of tap-events we calculated the intertap intervals (it is). However, for the comparison between empirical data and simulated data reported in the model validation section, we calculated and sampled the phases of the oscillators in steps of 10 ms for increased accuracy.

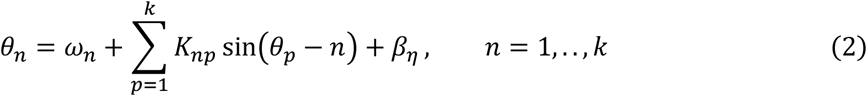

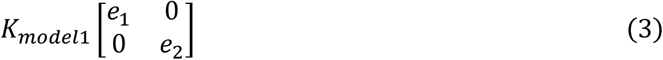

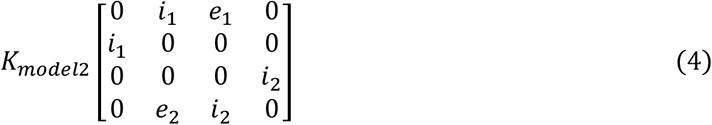

### Model behaviour

To determine the range of coupling weights wherein the models transitioned from an incoherent to coherent state we ran simulations over a wider range of coupling weights. We linearly increased coupling weights equally between the oscillators in both models, starting at a minimum coupling weight of 0.1 up to a maximum of 30 with a step size of 0.1. 200 simulations of 12 seconds each were averaged per step. Note that in model 2 we only considered synchronization between the two action oscillators. We found that the models quickly reached coherent synchronous state, as shown in Figure 5.

To examine the behaviour of the models we performed a parameter search within the previously determined bifurcation range. For the two-oscillator model we exploited the symmetrical nature of the two oscillators and coupling weights, by restricting the search range to a coupling weight of 0 to 5 for the between-unit coupling term 1 (*e*_1_) and 10 to 5 for between-unit coupling term 2 (*e*_2_) in steps of +− 0.1. This results in 51 possible combinations, and for each combination 200 simulations of 12 seconds were run. The oscillators were initiated at random phases, and the first two seconds of simulations were discarded to account for the metronome that was present in the first two seconds of the interaction in the empirical experiments. Four the four-oscillator model we had to restrict the search range, as a similar approach as with the two-oscillator model would results in over 5.7 × 10^8^ possible combinations. We decided to sample the model at four different coupling weights, 1, 5, 9, and 13. This gives 256 possible combinations, which we simulated in the same way as model 1.

We analysed the time series produced by the model by performing a cross-correlation at lag −1, 0 and +1 for each simulation run. These correlations coefficients were then averaged for each coupling weight combination. We then clustered the lagged cross-correlations using the complete linkage method, and performed a similarity profile analysis in R^38^, using the simprof-package^39^, at an adjusted alpha of 0.001 (shown in Figure 6)

**Figure 6.**
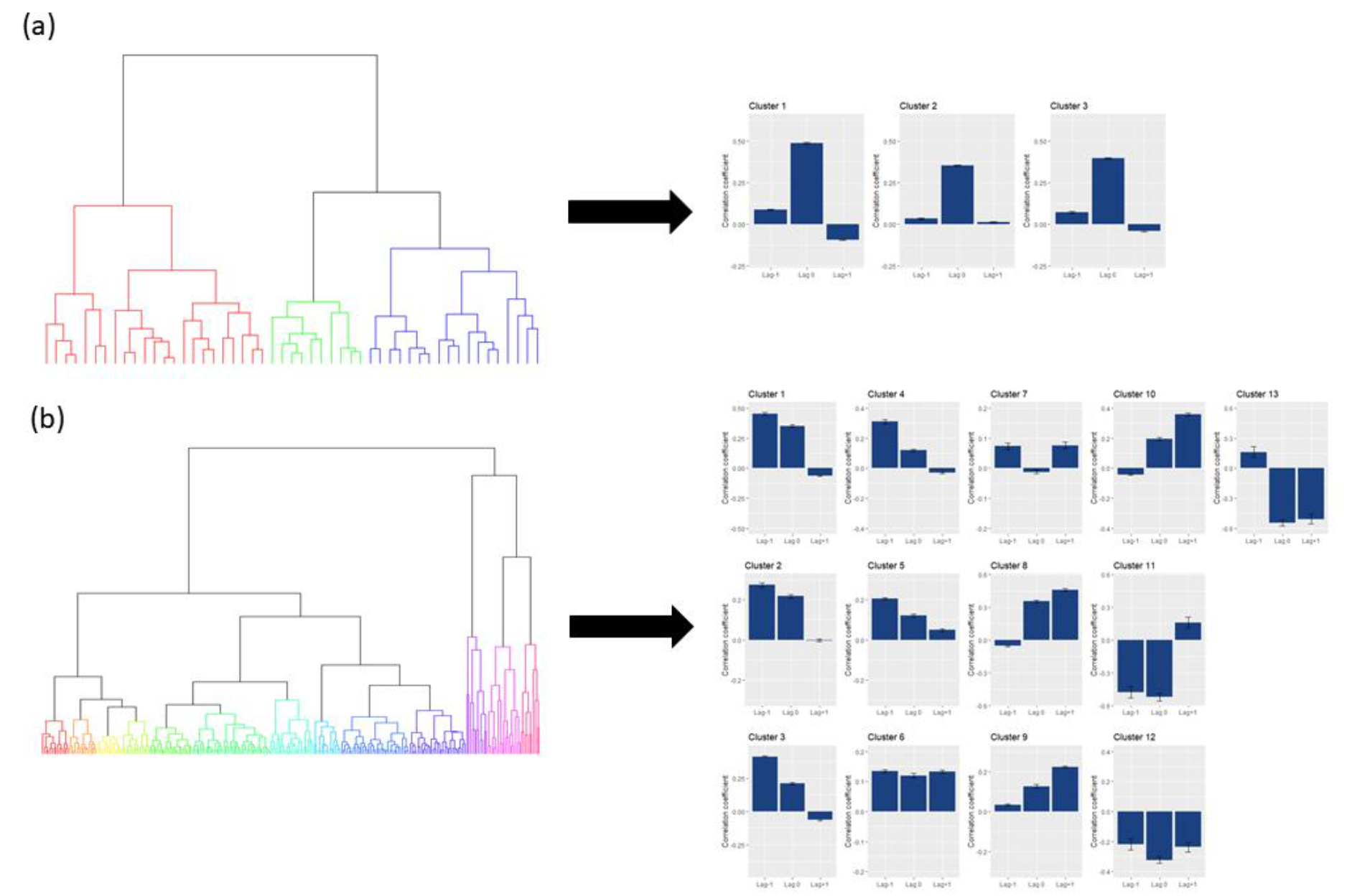
Clustering dendrogram and resulting lag patterns. In A the clustering dendrogram for a two-oscillator model is shown on the left. On the right, the corresponding mean lag patterns are shown. In this case, the two-oscillator model produced three significantly different patterns. All these three patterns were dominated by a strong lag-0 component. In B we show the same procedure applied to data from the four-oscillator model. Here we see a much richer variety of lag patterns, with 13 being significantly different.

### Model validation on empirical data

To calculate the coupling weight which best fitted to the empirical data we performed a two-step consecutive parameter search. For dataset 1 we searched for the coupling weights that resulted in the best fit with each of the three subgroups of participants in the empirical data. For each of the three separate subgroups we first simulated 300 trials for each possible coupling value combination between the four oscillators, with the coupling weights ranging from 1 to 15 in steps of 1. Each trial was simulated using the same approach as for study 1. We averaged the cross-correlated lags of the individual time series produced in each trial per coupling value combination, and calculated the numerical distance between the simulated data and the empirical data. The best fit was then selected, and a second search performed at coupling weights of +− 0.9 in steps of 0.2 following the same procedure. The resultant best fit based on numerical distance was then chosen, and its coupling weights were used to simulate 2000 trials. Here, we increased the accuracy in the time series data by calculating and sampling the phase of the oscillators in steps of 10 ms. The Bhattacharyya coefficient was calculated for each of the three lags (−1, 0, and +1) between the simulated data and the empirical data separately and then averaged, using the disparity package in R^40^. The same approach was followed for dataset 2. This approach resulted in approximately 8 × 10^11^ simulations, which were performed over the course of roughly 60 hours using a MATLAB Distributed Computing Server.

